# Phased Potato Genome Assembly and Association Genetics Enable Characterisation of the Elusive H1 Resistance Locus Against Potato Cyst Nematodes

**DOI:** 10.1101/2025.08.04.667887

**Authors:** Yuk Woon Cheung, Lynn H Brown, Thomas M Adams, Brian Harrower, Amanpreet Kaur, Gaynor McKenzie, Jamie Orr, James Price, Vikrant Singh, Moray Smith, Micha Bayer, Ingo Hein

## Abstract

The complexity of potato genetics, characterised by tetrasomic inheritance, has contributed to slower genetic gain in potato compared to other major crops. Disease resistance genes, often found in large clusters of highly similar paralogs and alleles, further complicate genetic studies. The *H1* resistance locus, introgressed into potato cultivars from *Solanum tuberosum* spp. *andigena*, has been successfully used for over 60 years to control *Globodera rostochiensis* in Europe. Although previous genetic studies mapped this resistance to chromosome 5, the complete structure of the locus remained elusive. To reduce genomic complexity, we generated a dihaploid of the cultivar ‘Athlete’, DH4_Athlete, carrying the *H1* resistance locus, and produced a phased haplotype representation of the *H1* interval using Oxford Nanopore sequencing. Combined with RenSeq-based association genetics, this approach allowed us to reconstruct the entire *H1* locus, including recombination points at both the 5′ and 3′ ends of the interval.

## Introduction

Potato (*Solanum tuberosum*) is the world’s most important non-cereal food crop, consumed by over a billion people and serving as a dietary staple in numerous countries (Birch et al., 2012; Wijesinha-Bettoni & Mouillé, 2019). In 2023, global production reached approximately 383 million tonnes, underlining its significance in global food and economic security (FAO, 2023). However, the sustainability of potato production is increasingly challenged by biotic stresses, particularly pests and pathogens, which collectively account for an estimated 21% yield loss (Savary et al., 2019). Among these, potato cyst nematodes (PCN) pose one of the most important threats, contributing to around 9% of total yield reductions worldwide whilst threatening the potato supply chain as contaminated land is unsuitable for seed potato production (Gartner et al., 2021).

PCN in Europe commonly refers to two major species, *Globodera rostochiensis* and *G. pallida*, soil-borne nematodes with complex, obligate parasitic life cycles. A major challenge in PCN management is their ability to remain dormant in soil, enabled by a tightly regulated hatching process. After fertilisation, the cuticle of the gravid female hardens, forming a cyst that encases often hundreds of eggs. This cyst detaches from the host roots and persists in the soil, with eggs hatching only in response to specific chemical signals released by the roots of a suitable host plant (Price et al., 2021). This remarkable longevity makes PCN difficult to eradicate and has led to its designation as a quarantine pest in over 100 countries, severely restricting international seed potato trade and causing significant economic losses (Bairwa et al., 2024).

Resistance breeding against *G. rostochiensis* has yielded notable successes with the discovery of the *H1* resistance, which confers near-complete resistance to the Ro1 and Ro4 pathotypes that are prevalent in many potato growing countries outside South America (Evans & Brodie., 1980). The resistance phenotype was first identified in the wild subspecies *Solanum tuberosum* ssp. *andigena* CPC 1673, collected by Dr. Cárdenas in La Paz, Bolivia, and received in the UK in December 1944. Notably, the material arrived as insect-damaged tubers, which were ultimately discarded by 1949. The line survived only as true seed, which was later used in a sibling cross to produce TBRADG 3520 in 1962 (G. McKenzie, personal communication).

Although the *H1* gene has not yet been cloned, it has been mapped to the distal end of chromosome 5, a region rich in NLR (nucleotide binding leucine rich-repeat) resistance genes. This region is difficult to resolve due to low recombination rates and high haplotypic diversity. Previous studies, including those by Bakker et al., (2004) and Finkers-Tomczak et al., (2011) defined a region within the resistant haplotype SH83-92-488. However, this region did not reveal the causal gene responsible for resistance to *G. rostochiensis*.

NLR proteins are a major class of intracellular immune receptors in plants, typically comprising a central NB-ARC (nucleotide-binding adaptor shared by APAF-1, certain *R* gene products, and CED-4) domain, a C-terminal LRR (leucine-rich repeat) domain, and an N-terminal CC (coiled-coil) or TIR (toll/interleukin-1 receptor-type) domain. To target and analyse these genes, RenSeq (Resistance gene enrichment Sequencing) was developed to specifically sequence NLR-encoding genes from complex plant genomes (Jupe et al., 2013). This approach has enabled comprehensive annotation and evolutionary studies of the NLRome, the complete set of NLR genes in a genome.

RenSeq has since evolved into a suite of methods, including SMRT-RenSeq (PacBio-based long-read sequencing) (Witek et al., 2016), AgRenSeq (*k*-mer based association genetics) (Arora et al., 2019), and dRenSeq (diagnostic detection of known NLRs) (Armstrong et al., 2019; Van Weymers et al., 2016). These were recently unified into the SMRT-AgRenSeq-d pipeline, which has proven effective in resolving complex loci, such as the *Gpa5* resistance gene in tetraploid potato (Adams et al., 2025; Wang et al., 2023).

In this study, we initially used SMRT-AgRenSeq-d to identify candidate NLR genes associated with *H1*-mediated resistance in the cultivar ‘Buster’. To develop a complete and accurate representation of the *H1* locus, we adapted AgRenSeq to produce phased Oxford Nanopore haplotypes from the dihaploid clone DH4_Athlete, derived from the resistant cultivar ‘Athlete’. We further demonstrate that the resolved haplotype alone represents a significant advancement for genome-based association genetics, enabling high-resolution analysis of trait associations. This combination of phased haplotypes and association genetics provides a detailed representation of the *H1* haplotype, including recombination events at both the 5′ and 3′ ends of the locus.

## Results

### SMRT-RenSeq

The potato cultivar ‘Buster’, released in 2020, is well known for its robust resistance to potato cyst nematode (PCN), particularly *G. rostochiensis* pathotypes Ro1 and Ro4, and was used as a reference to represent the NLRome of a resistant cultivar. Using the SMRT-RenSeq workflow from the nfHISS pipeline (Adams et al., 2025; Wang et al., 2023), contigs were assembled from PacBio HiFi RenSeq reads of ‘Buster’. The assembly yielded 6,292 contigs with an N50 of 11,379 bp. Of these, 2,017 contigs contained a total of 2,870 NLR-like sequences, of which 1,537 were annotated as complete NLRs, with the remainder made up of pseudogenous and/or incomplete NLRs. A dRenSeq analysis confirmed the presence of several previously cloned resistance genes in ‘Buster’, including *Rpi-R3a, Rpi-R3b, Rx*, and *Gpa2*. Each of these genes were fully and accurately represented on individual contigs in the PacBio HiFi assembly (Table 1).

**Table 1.** Representation of four benchmark genes following HiFi-based assembly.

| Buster Reference NLR representation |  |  |  |  |  |  |  |  |  |
| --- | --- | --- | --- | --- | --- | --- | --- | --- | --- |
| Referen<br>ce<br>Gene | Representat<br>ive<br>Sequence | Identi<br>ty (%) | Gen<br>e<br>lengt<br>h | Mismat<br>ch | Ga<br>p | Referen<br>ce gene<br>start | Referen<br>ce gene<br>end | Covera<br>ge of<br>gene<br>(%) | Coverag<br>e of HSF<br>(%) |
| Rpi_R3<br>a | tig00000259 | 100 | 3849 | 0 | 0 | 1 | 3849 | 100 | 100 |
| Rpi_R3<br>b | tig00000358 | 100 | 3852 | 0 | 0 | 1 | 3852 | 100 | 100 |
| Gpa2 | tig00002324 | 100 | 2976 | 0 | 0 | 1 | 2976 | 100 | 100 |
| Rx | tig00001251 | 100 | 3045 | 0 | 0 | 1 | 3045 | 100 | 100 |

### Development of an association panel

To investigate the genetic basis of the H1 resistance and to identify candidate resistance genes at the *H1* locus, we developed an association panel comprising 126 potato cultivars. This panel was carefully curated to include accessions with contrasting phenotypes: 58 highly resistant and 68 highly susceptible cultivars. Phenotypic data were compiled from a combination of publicly available databases and commercial sources. A detailed summary of the selected cultivars and their resistance profiles is provided in Table S1. Notably, the panel includes TBRADG 3520 (CPC 3520), a resistant sibling clone of CPC 1673, the original source of *H1*-mediated resistance.

### SMRT-AgRenSeq-d

The SMRT-AgRenSeq-d analysis, applied to the association panel described above (Table S1), identified candidate NLR contigs associated with *H1*-mediated resistance in the cultivar ‘Buster’. The association mapping, visualised in Figure 1A, revealed 89 contigs which contained 131 NLRs with varying association scores based on a *k*-mer presence/absence matrix. These associated contigs, marked in red, were identified using an inclusive threshold of half the maximum association score for first pass candidate identification.

**Figure 1.**
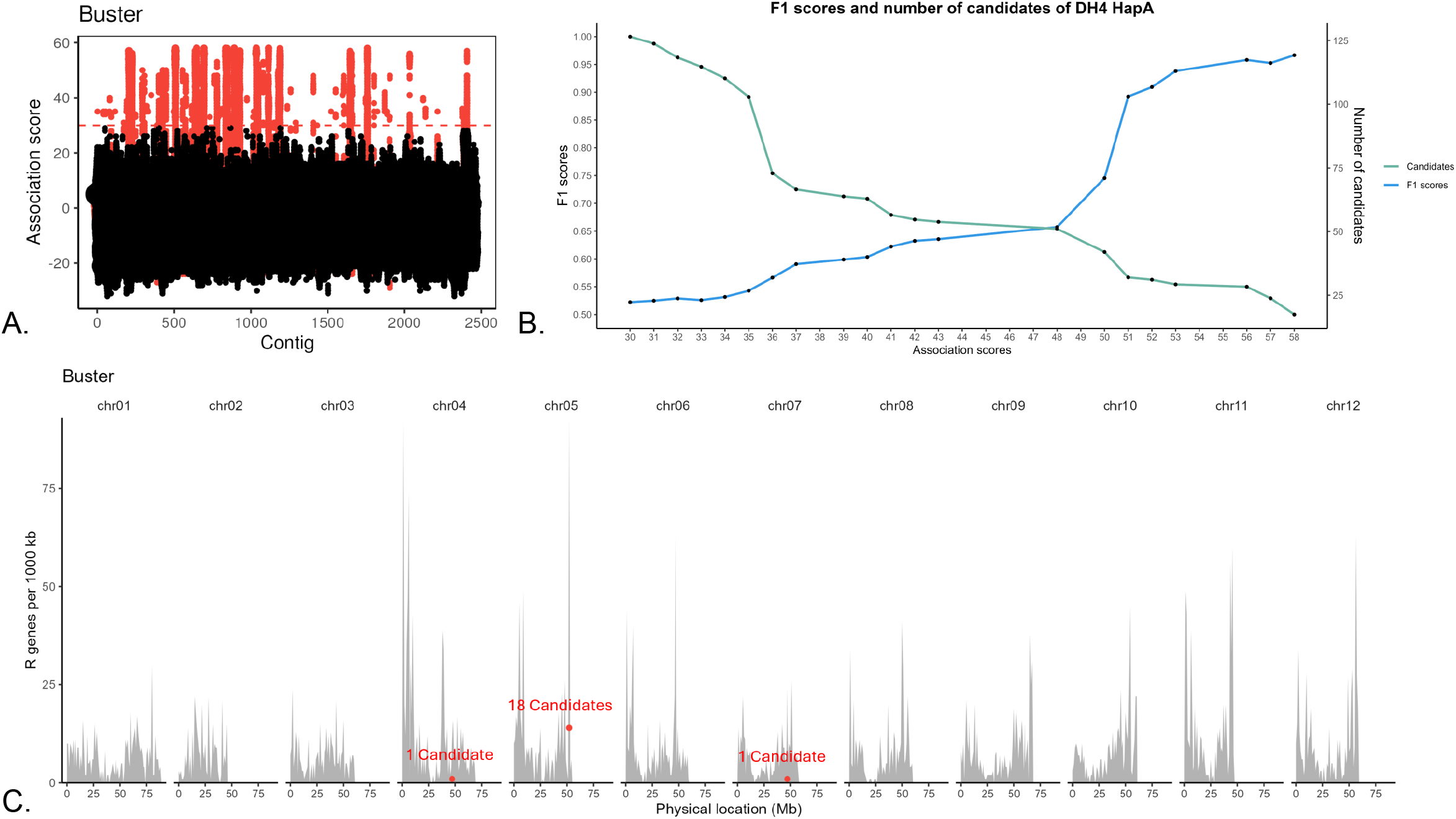
SMRT-AgRenSeq-d analysis of H1. (A) identification of NLR containing contigs in the reference cultivar ‘Buster’ with association to H1. Columns on the x-axis represent NLR containing contigs, with dots indicating mapped k-mers and their association score to H1. The dashed line highlights the chosen threshold (50% of the maximum = 29) for inclusive candidate selection. (B) F1 scores and number of candidate NLRs for each association score group. (C) The predicted positions of H1 candidates with F1 score >0.95 are shown (red dots) in relation to all NLRs from reference Buster mapped onto DM v6.1 over all 12 chromosomes.

To refine candidate selection, dRenSeq was used to determine the presence or absence of NLR contigs across the resistant and susceptible cultivars, allowing for the calculation of F1 scores (Figure 1B; Table S2). The F1 score, where absolute linkage is denoted by 1 and any recombination leads to a score of less than 1, identified a subset of 20 NLR contigs surpassing an F1 score of 0.95, of which 5 candidates had an F1 score of 1, confirming that candidates with higher association scores consistently exhibited increased predictive accuracy, indicating strong genotype-phenotype concordance. Mapping these high-confidence candidates onto the reference genome of the susceptible clone DM v6.1 (Figure 1C) shows their predominant localisation to the distal end of chromosome 5, aligning with the previously described genomic position of the *H1* resistance locus. However, two candidates map onto chromosomes 4 and 7 of DM v6.1 respectively.

To further investigate the relationships among the highly associated and genetically linked candidate NLRs, a phylogenetic tree was constructed using the NB domains from SMRT-RenSeq-derived NLRs in ‘Buster’ (Figure 2). The resulting tree revealed that all high-confidence candidates clustered tightly within a single clade, suggesting a common evolutionary origin and, potentially, functional relatedness. This clade also includes known resistance genes such as *Mi-1.1* and *Mi-1.2*, two wild tomato (*Solanum peruvianum*) genes that confer resistance to root-knot nematodes and several phloem-feeding insects (Rossi et al., 1998), as well as *Rpi-blb2*, an NLR from *Solanum bulbocastanum* that provides resistance to *Phytophthora infestans* (van der Vossen et al., 2005). These findings are consistent with earlier studies of the *H1* locus, which identified homologous resistance genes with approximately 52% identity to *Rpi-blb2*, a member of the CC-NB-LRR class of resistance proteins (Finkers-Tomczak et al., 2011). One exception was the candidate on tig0000215, which lacked an identifiable NB domain and was therefore excluded from the analysis.

**Figure 2.**
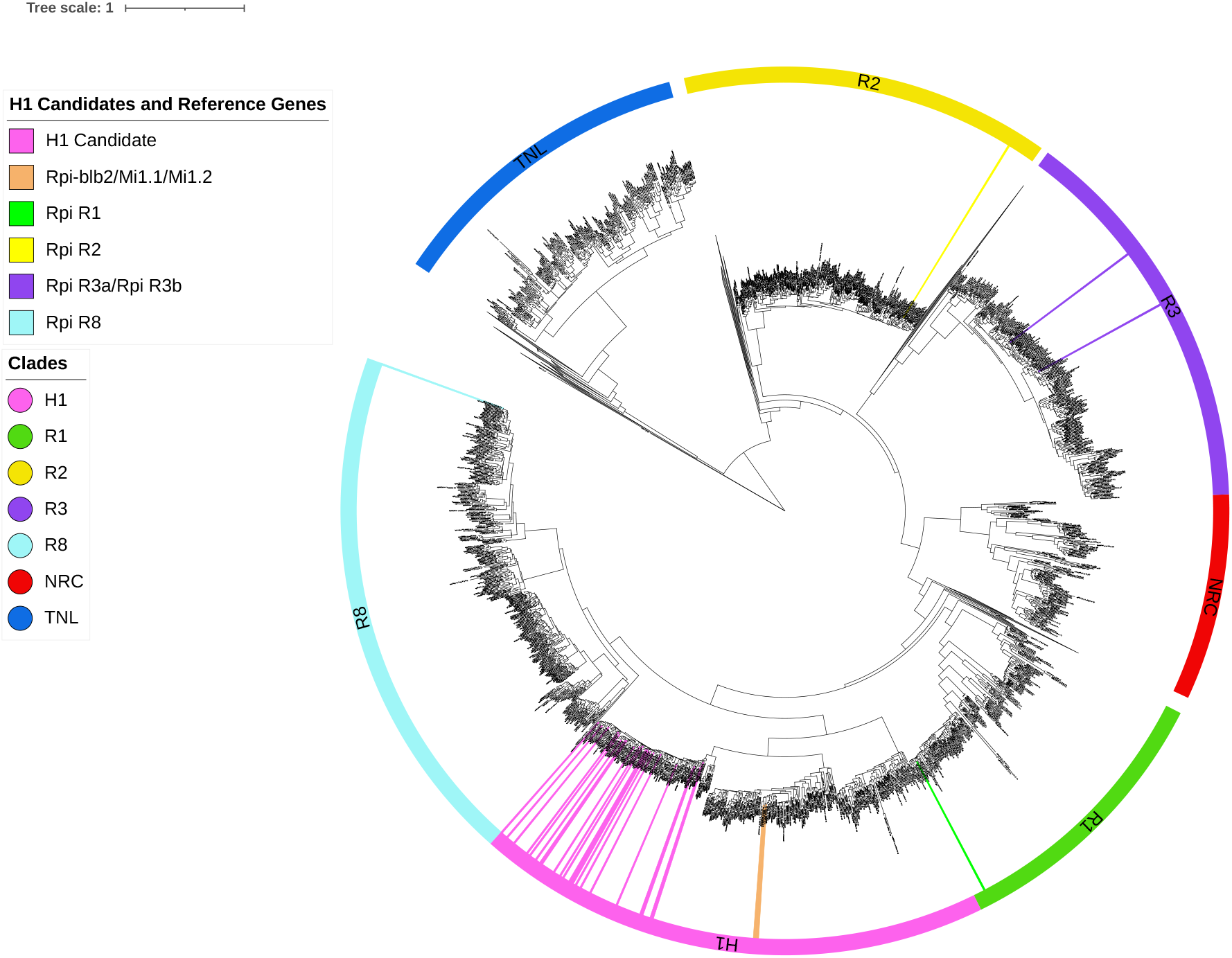
A. Phylogenetic tree displaying NB domains of ‘Buster’ NLRs following assembly of SMRT-RenSeq reads. Expanded are the clades containing previously identified functional NLRs such as *Rpi-R1* (highlighted in green), *Rpi-R2* (Highlighted yellow), *Rpi-R3a* and *Rpi-R3b* (highlighted in purple) alongside NRCs from tomato (*Solanum lycopersicum*), *Nicotiana benthamiana* and *Beta vulgaris*. Collapsed clades consist of NLR clades which are not investigated in this study. Highlighted in pink are the H1 candidates with F1 score > 0.95 (No NB domain was identified in the candidate on tig00002155, hence it was excluded from this analysis). The NB containing proteins *APAF1* from *Homo sapiens* and *CED4* from *Caenorhabditis elegans* act as outgroups. The tree scale indicates substitutions/site. B. Zoomed-in view of candidate H1 clade. Highlighted in pink are H1 candidates with F1 score > 0.95. Highlighted in orange are reference genes *Mi-1.1* and *Mi-1.2* from *Solanum peruvianum*, and *Rpi-blb2* from *Solanum bulbocastanum*.

### ONT Sequencing of a Dihaploid derived from ‘Athlete’

Since the DM v6.1 reference genome lacks *H1*-mediated resistance, it can only serve as a proxy for genomic studies at this locus. To address this limitation, a physical representation of the H1 locus was generated using long-read Oxford Nanopore Technologies (ONT) sequencing of a dihaploid clone derived from the cultivar ‘Athlete’ (DH4_Athlete). This assembly captured all 20 previously identified H1 candidate NLR genes with F1 scores > 0.95, along with four additional candidates with F1 scores < 0.95 (Table S3). Furthermore, the feasibility of using a haplotype-resolved genome assembly directly for association mapping was explored, with the goal of eliminating the need for separate PacBio-based NLRome assemblies.

ONT sequencing of DH4_Athlete yielded 13.9 million reads, with an N50 of 23.5 kbp and 77.8% of reads above Q20, providing approximately 84-fold coverage based on the reference potato genome DM v6.1 size of 840 Mb (The Potato Genome Sequencing Consortium., 2011). *K*-mer spectrum analysis confirmed the genome to be diploid (Figure S1). A partially phased assembly representing both haplotypes was produced. Haplotype A consisted of 1,892 contigs (849 Mb total, N50 = 57.3 Mb), while Haplotype B comprised 100 contigs (779 Mb, N50 = 31.3 Mb). Both haplotypes showed high completeness (BUSCO > 96%), with 710 NLRs in Haplotype A and 646 in Haplotype B. Both haplotypes exhibited a high degree of gene synteny and contiguity, with multiple chromosomes being resolved as telomere-to-telomere (Table 2, see also Figures S2 and S3).

**Table 2.** Assembly statistics for the dihaploid, partially phased DH4_Athlete genome based on ONT sequencing data.

|  | Haplotype A | Haplotype B |
| --- | --- | --- |
| **Contig N50** | 57.3Mbp | 31.3Mbp |
| **# Sequences** | 1,892 | 100 |
| **Total assembly size** | 849Mbp | 779Mbp |
| **BUSCO (Single; Duplicated)** | 96.1% (90.7%; 5.4%) | 96.9% (93.3%; 3.6%) |
| **# NLRs** | 710 | 646 |
| **GCI score** | 36.7 | 24.5 |
| **GCI contig #** | 45 | 46 |

### Identification of NLRs in DH4_Athlete

Genes were predicted *de novo* with Helixer (Holst et al., 2023), from which NLRs were identified with Resistify (Martin et al., 2022; Smith et al., 2025). As Helixer retrieved a greater number of canonical NLRs across the phased assembly, and manual inspection of NLR gene models indicated both tools performed similarly, Helixer-predicted NLRs were taken forward for analysis. This yielded a total of 1,356 NLRs across both haplotypes, with approximately 10 percent more NLRs represented in haplotype A (Table 2).

To validate the candidate genes identified in ‘Buster’ through SMRT-AgRenSeq-d, the partially phased haplotype assembly of DH4_Athlete was utilised for an association study using the same panel of varieties with contrasting phenotypes (Table S1). This analysis demonstrated that strong association peaks existed in both Haplotype A (Figure 3A) and Haplotype B (Figure 3B). A dRenSeq analysis only identified candidates with an F1-score of 1 in Haplotype A and not in Haplotype B (Figure 3C; Tables S4 and S5).

**Figure 3.**
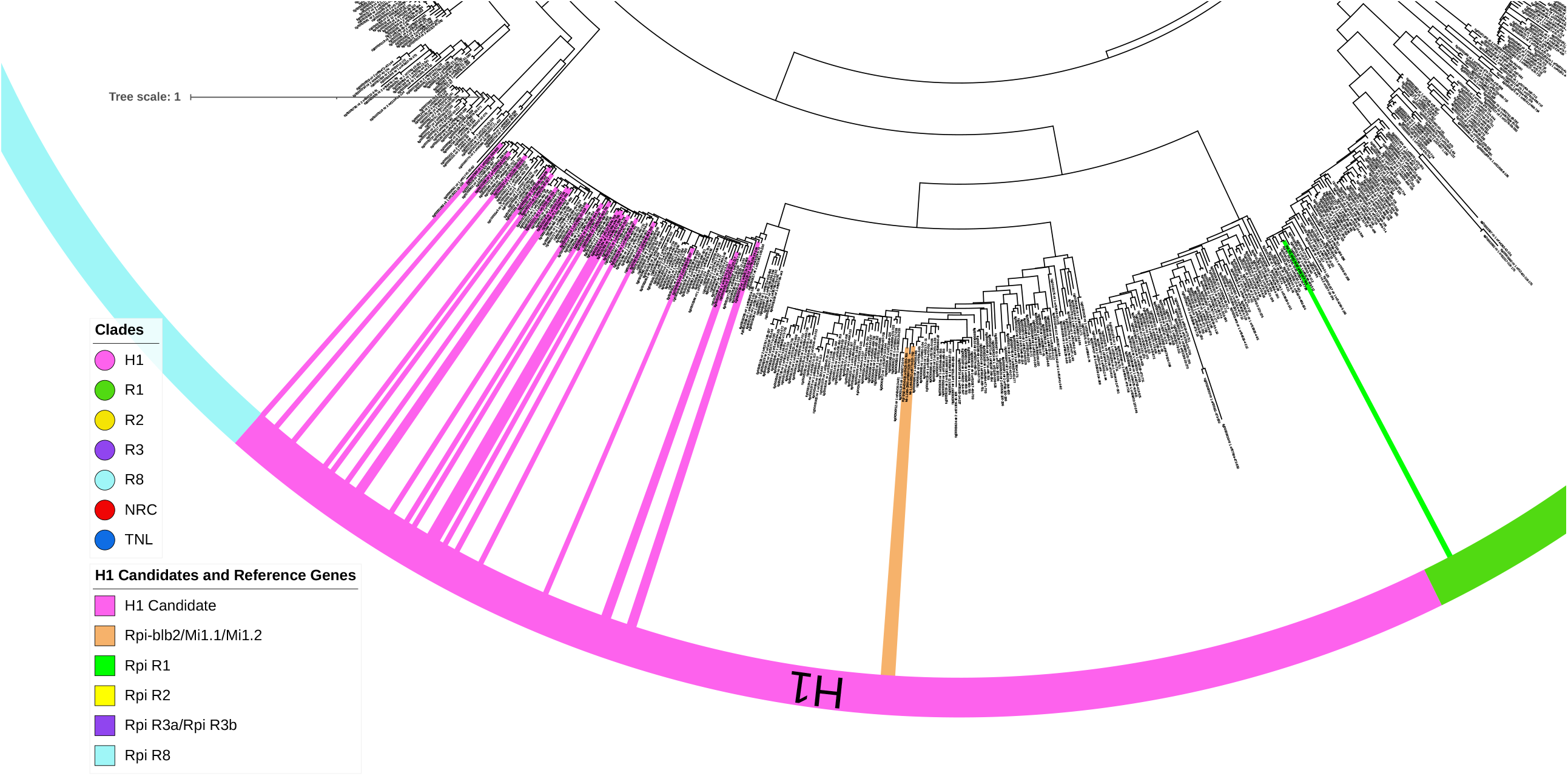
A. Association plot showing NLRs extracted from DH4_Athlete haplotype A with mapped k-mers and their respective association scores for H1. The x-axis represents individual NLRs, while dots on the y-axis indicate k-mer associations. The dashed red line marks the chosen association threshold (score = 29, e.g., 50% of maximum). B. Equivalent association plot for DH4_Athlete haplotype B. C. Boxplot of F1 scores for the 20 and 10 H1 candidate NLRs identified in the association studies with reference DH4_Athlete haplotype A and B, respectively.

Further, analysis of *k*-mer counts across candidates from both haplotypes revealed that the strongest association is found in the partially phased haplotype A, rather than haplotype B (Figure S4). This suggests that NLRs in haplotype B share some limited conserved regions with *H1* candidates. Supporting this, a BLAST comparison confirmed that candidate athlete_chr05_B_001411 from haplotype B shares 98% identity covering 82% of the candidate athlete_chr05_A_001451 from haplotype A, accounting almost exclusively for the elevated *k*-mer count and association signal (Table S6).

*H1* candidates identified using ‘Buster’ in the PacBio-based SMRT-AgRenSeq-d pipeline showed strong concordance with those from the ONT-based DH4_Athlete association study, as confirmed by BLAST (Table S7). Minor differences mainly arose from how coding sequences (CDS) were defined. The ONT-based pipeline used *Helixer* for *de novo* gene prediction and full gene model annotation, followed by NLR detection with *Resistify*. In contrast, the SMRT-AgRenSeq workflow used *NLR-Annotator*, which identifies NLR motifs but not full gene structures, instead inferring CDS boundaries based on the inclusion of 1 kb flanking regions, sometimes leading to truncated predictions.

For example, a ‘Buster’-based candidate on tig00002155 lacked an NB domain and was absent in ONT predictions, as *Resistify* only reports NB-ARC-containing genes. ONT-based annotations also tended to predict longer genes. One ONT gene, DH4_Athlete_Chr05_A_001462, was 16 kbp longer than its *Buster* counterpart, contributing to a lower F1-score (< 0.95), likely due to bait coverage bias. Despite such discrepancies, high-confidence candidates (F1 > 0.95) were closely clustered, and those with absolute linkage (F1 = 1) fell within a ~270 kbp interval. Lower-scoring candidates (F1 < 1) in the same region reflect both structural and coverage-related variation.

Importantly, despite differences in gene structure predictions, all high-confidence candidate genes and their corresponding contigs identified by SMRT-AgRenSeq-d in ‘Buster’ were independently mapped to Haplotype A of DH4_Athlete (Figure 4). Notably, two high-confidence candidates initially mapped to chromosomes 4 (tig00002170_nlr_1) and 7 (tig00001229_nlr_1) of the DM v6.1 reference (Figure 1C) were reliably located within the H1 cassette on chromosome 5 in DH4_Athlete Haplotype A, represented by athlete_chr05_A_001138 and athlete_chr05_A_001460 (Figure 4; Table S7).

**Figure 4.**
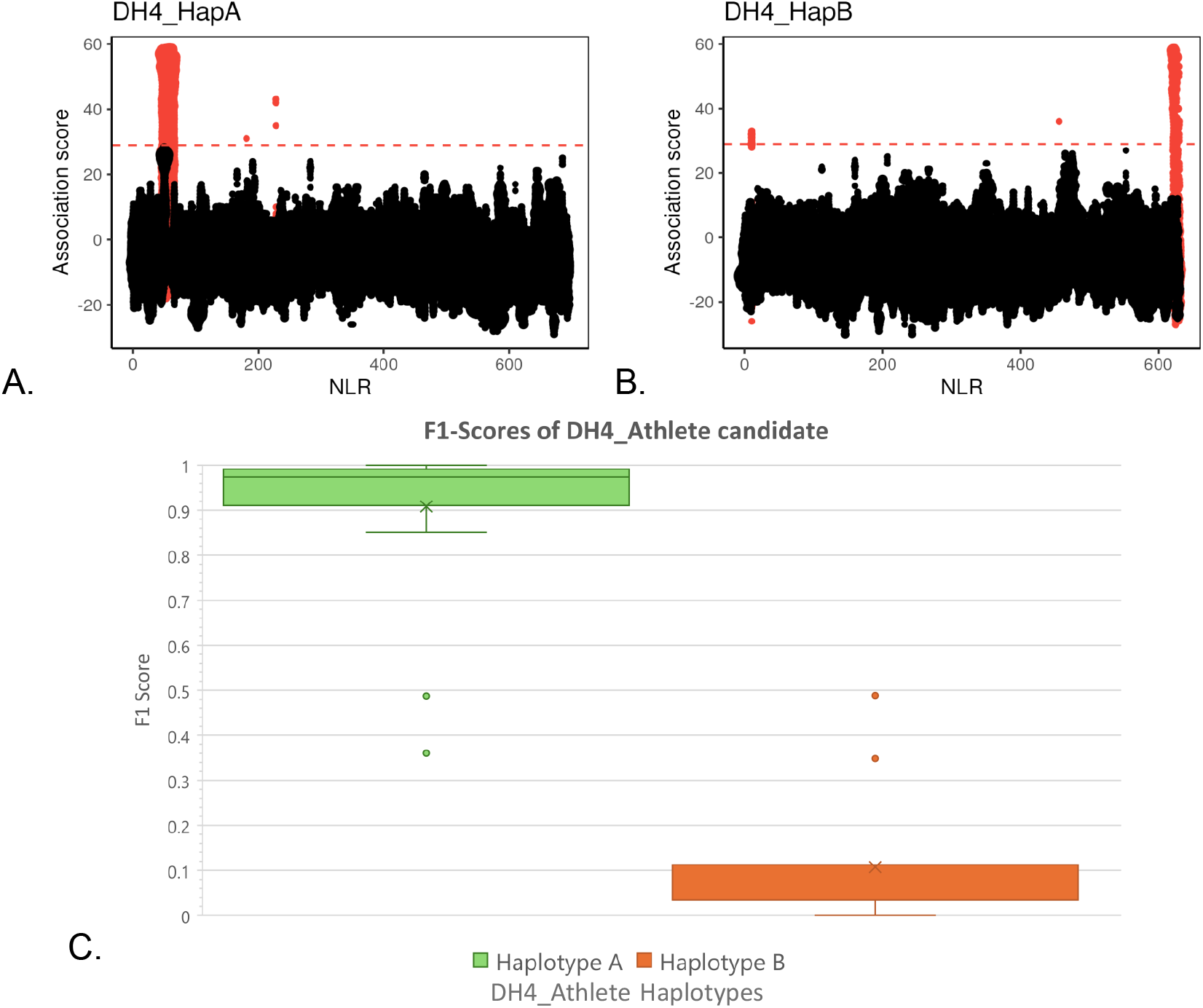
Representation of predicted candidate genes in reference to DH4_Athlete Haplotype A. The upper panel shows the DH4_Athlete haplotype A of chromosome 5, with the H1 locus highlighted in red at the distal end (~54Mb). Below, the predicted candidate genes from the association study using DH4 Athlete as a reference within this region are represented. The lower panel illustrates the conservation of the H1 cassette across various resistant (green bars) and susceptible (orange bar) cultivars. Candidates that are identical to SH genes previously reported by Finkers-Tomczak et al. (2011) are also indicated. Below are the candidate genes from the cv. ‘Buster’ SMRT-AgRenSeq-d assembly that mapped within this cassette.

As shown in this study, ONT-based sequencing offers key advantages for NLR analysis, notably its ability to generate long reads that span complex genomic regions, including full-length NLR genes, structural variants, and potentially multiple NLRs within gene clusters (Figure 5). This facilitates accurate physical mapping of resistance loci and reveals gene clusters that are often fragmented or collapsed in PacBio HiFi and short-read assemblies. ONT is especially effective for large, repetitive genomes like tetraploid potato, or, as shown here, a dihaploid of ‘Athlete’ (DH4_Athlete) with reduced complexity. However, current ONT assemblies still face challenges in highly heterozygous polyploid genomes, where phasing is limited and closely related alleles may be collapsed. By integrating ONT sequencing with AgRenSeq-d, many of these limitations are addressed. This combined approach enables k-mer-based association to contigs, supporting haplotype-specific analysis and more precise allele resolution.

**Figure 5.**
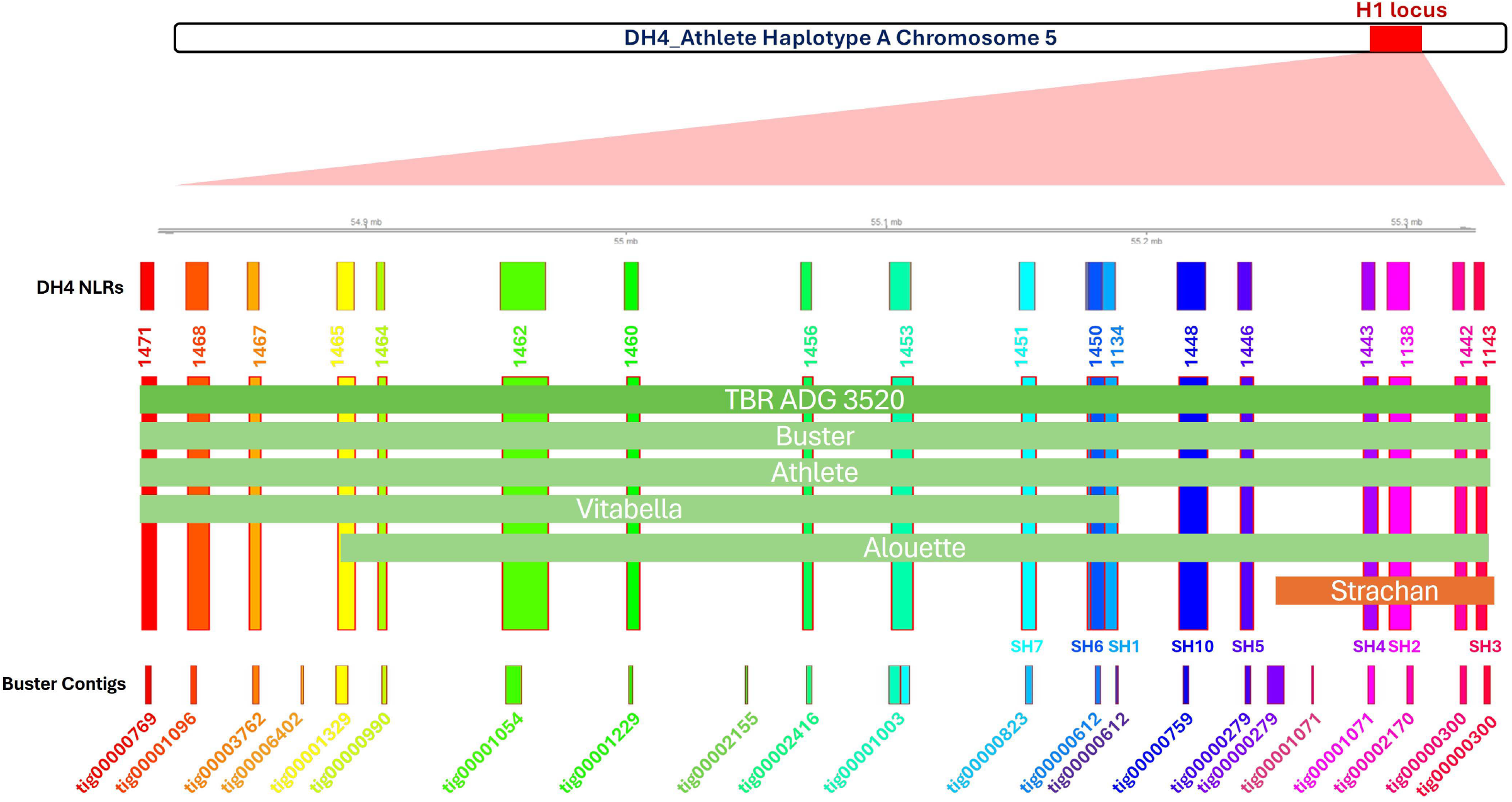
Gene synteny at the H1 locus, defined by outermost genes of the haplotype A cluster.

In this study, ONT sequencing allowed us to fully resolve the H1 locus, including recombination boundaries and introgressed gene clusters. Importantly, the phased assembly of DH4_Athlete revealed that all high-confidence candidates from association genetics were located exclusively in haplotype A, highlighting the value of haplotype-specific information for distinguishing causal genes from non-functional homologues (Figures 4 and 5). These findings represent the first genetically validated physical map of the H1 resistance locus in potato.

The dRenSeq analysis confirmed that the introgression cassette from the resistant CPC 1632 sibling clone, the source of the original *H1* introgression, remains highly conserved. Finkers-Tomczak et al., (2011) identified 17 resistance gene homologs (RGHs) spanning ~314 kb within the *H1* cassette of the diploid clone SH83-92-488 (SH), including genes SH1 through SH10. In our analysis, we detected SH1–SH7 and SH10, indicating strong conservation in the northern region of the interval, previously linked to *H1* resistance in SH. However, expanded analysis across additional varieties revealed recombination events within SH2, SH3, SH4, SH5, and SH10. These genes were absent in ‘Vitabella’ and ‘Alouette’ but present in ‘Strachan’, a susceptible cultivar, suggesting recombination has decoupled resistance from these genes. We also observed variation in SH8 and SH9. For example, these genes are present in ‘Vitabella’, but absent in TBRADG 3520, ‘Buster’, and DH4_Athlete (confirmed by dRenSeq) (Table S8), helping to pinpoint a recombination breakpoint that separates the conserved region of the cassette (Figure 4).

Using the presence and absence of candidates that are highlighted in the cultivars ‘Strachan’, ‘Vitabella’ and ‘Alouette’, the conserved *H1* introgression cassette can be narrowed down to approximately 270 kilobases in the H1-containing haplotype A, which is absolutely linked to the *H1* resistance phenotype. A comparison of the corresponding loci of DM v6.1 as well as the susceptible haplotype B provides evidence of significant locus expansion and rearrangement of NLRs in haplotype A (Figure 5).

In summary, these results highlight the value of integrating long-read sequencing, partially phased genome assemblies, and association genetics to dissect complex resistance loci. This framework enables more effective candidate gene identification and supports informed, precise breeding strategies, not only for PCN resistance, but also for a broad range of traits governed by complex genomic architecture in polyploid crops.

## Materials and Methods

### Extraction of DNA

As described in Wang et al., (2023), all potato accessions’ DNA was extracted using the Qiagen DNeasy Plant Mini Kit (Qiagen, Hilden, Germany) according to manufacturer’s instructions. DNA from the cultivar ‘Buster’ for HiFi RenSeq sequencing was extracted according to manufacturer’s instructions for the Wizard ® HMW DNA extraction Kit (Promega, Madison, WI, USA). High-molecular-weight DNA was extracted from fresh leaf tissue of dihaploid plants of the tetraploid potato cultivar ‘Athlete’ DH4_Athlete after three days of dark treatment, following the manufacturer’s guidelines for the NucleoBond HMW DNA kit (Macherey-Nagel, Germany).

### RenSeq

The RenSeq bait library version 5 was used as described in Armstrong et al., (2019). Enrichment sequencing was performed by Arbor Biosciences (Ann Arbor, MI, USA) using the Novaseq 6000 platform for Illumina reads and the PacBio Sequel II platform for HiFi reads.

### Selection of Association Panel

The identification of *H1* used a panel of 126 cultivars which consisted of highly resistant and highly susceptible accessions determined by retrieving results from various databases and sources. Within the association panel were 58 resistant and 68 susceptible varieties. Phenotypes of these varieties were collected from various sources including: the European cultivated potato database, the AHDB potato database, Europlant, Solanum International Inc., Potato Pro, Bavaria Saat, The Potato Company, Greenvale, Solana United Kingdom, and from a previous study by Schultz et al., (2012) which has marker presence data available, summarised in Table S1.

Computational analyses were performed on the James Hutton Crop Diversity HPC (High Performance Compute cluster), described by Percival-Alwyn et al., (2025).

### SMRT-AgRenSeq-d

The nfHISS workflow version v1.1.0 was used to perform SMRT-AgRenSeq-d which includes SMRT-RenSeq assembly, AgRenSeq and dRenSeq as described in Adams et al. (2025) and validated in Wang et al., (2023).

### ONT-AgRenSeq-d

DH4_Athlete is a dihaploid plant of the tetraploid potato cultivar ‘Athlete’, generated by crossing with IVP (in vitro pollinator) line 48. ONT sequencing libraries were prepared using the Ligation Sequencing gDNA – Native Barcoding Kit 24 V14 kit, according to manufacturer’s guidelines. Libraries were sequenced on a combination of minION and PromethION R10.4.1 flow cells. After 24 hours, flow cells were washed and reloaded with the same library, following the guidelines of the Flow Cell Wash Kit (EXP-WSH004; Oxford Nanopore Technologies).

Reads were basecalled using Dorado v0.9.6 (Kuśmirek, 2023) with the dna_r10.4.1_e8.2_400bps_sup v4.1.0 and v5.1.0 models, depending on sampling rate. Genome ploidy and heterozygosity was determined with genomescope2 v2.0.1 (Ranallo-Benavidez et al., 2020), with 21-mers counted from ONT reads with kmc v3.2.4 (Kokot et al., 2017). Partially phased contigs were produced using hifiasm v0.25.0 (Cheng et al., 2021) with the --ont model and independently scaffolded to the doubled monoploid potato DM 1-3 516 R44 (v6.1) genome. Unscaffolded contigs were removed, and the resulting chromosomal scaffolds were merged into a single file. Genome completeness was evaluated with BUSCO v5.8.3 (Manni et al., 2021), using miniprot (Li., 2023) with the solanales_odb12 lineage. The contiguity of each haplotype assembly was assessed with GCI v1.0 (Chen et al., 2024), aligning ONT reads with minimap2 v2.30 (Li., 2018) and winnowmap v2.03 (Jain et al., 2022), with 15-mers counted by meryl v1.4.1 (Rhie et al., 2020).

Genes were predicted *ab initio* with Helixer v0.3.4 (Holst et al., 2023) using the model land_plant_v0.3_a_0080, and transposable elements with HiTE v3.3.2 (Hu et al., 2024). NLRs were identified and classified from the gene annotations with Resistify v1.2.0 (Martin et al., 2022; Smith et al., 2025) with CoCoNat (Madeo et al., 2023) enabled to increase CC sensitivity. The resulting NLR predictions were then passed through the AgRenSeq workflow.

### F1 Score

Candidate genes extracted from our highly associated contigs were further refined through generating F1 scores as described by Wang et al. (2023) and Chinchor (1992).

An F1 score of 1 means the classification achieved perfect precision and recall, correctly identifying all varieties as resistant or susceptible without any errors. The F1 score allows us to identify candidate genes reliably and accurately based on dRenSeq coverage.

### Phylogenetics

The phylogenetics workflow was as described by Wang et al. (2023). Briefly, NB domains were identified using InterProScan version 5.54-87.0 (Jones et al., 2014). The NB sequences were aligned with Clustal Omega version 1.2.4 (Sievers et al., 2011). Phylogenetic trees were constructed in R version 4.1.2 using the ape version 5.8-1 (Paradis & Schliep, 2019) and phangorn version 2.12.1 (Schliep et al., 2008) packages, applying the JTT+G substitution model, identified as the most appropriate model through BIC score, and 1,000 bootstrap replicates after alignment quality control. Clades were assigned using a 0.05 percentile threshold with PhyloPart version 2.1 (Prosperi et al., 2011), and the tree was visualised using IToL version 6.6 (Letunic & Bork, 2021).

## Data Availability

Raw reads are available at the European Nucleotide Archive (https://www.ebi.ac.uk/ena/browser/home) under accession numbers PRJEB89657 and PRJEB56823. Assemblies are available under the accession numbers PRJEB90064 and PRJEB90070.

## Code Availability

Scripts for ONT assembly of DH4_Athlete are available at Zenodo under URL https://doi.org/10.5281/zenodo.16418965.

## Acknowledgements

This work was supported by the Rural & Environment Science & Analytical Services (RESAS) Division of the Scottish Government through project JHI-B1-1, the Biotechnology and Biological Sciences Research Council (BBSRC) through award BB/X009068/1 and the Innovate UK Project 10058434: TRIP sustainable Agriculture. YWC was supported through the BBSRC PhD training program CTP-SAI award number BB/Y513684/1. LB was supported through the East of Scotland Bioscience Doctoral Training Partnership (EASTBIO DTP), funded by the BBSRC award BB/T00875X/1. JP, TA, MB and IH were supported through the Scottish Plant Health Centre project “PCN Action Scotland”.

The authors acknowledge Research Computing at the James Hutton Institute for providing computational resources and technical support for the “UK’s Crop Diversity Bioinformatics HPC” (BBSRC grants BB/S019669/1 and BB/X019683/1), use of which has contributed to the results reported within this paper.

## Supplemental Information

Table S1. Phenotypic data of varieties used in the Association studies.

Table S2. dRenSeq of 132 H1 candidates from SMRT-RenSeq reference ‘Buster’ in 126 cultivars

Table S3. BLAST results of all the candidates which map at 100% onto DH4_Athlete Chromosome 5 Haplotype A

Table S4. dRenSeq of 20 H1 candidates from ONT reference DH4_Athlete Haplotype A in 126 cultivars

Table S5. dRenSeq of 10 H1 candidates from ONT reference DH4_Athlete Haplotype B in 126 cultivars

Table S6. BLAST Top hit results of candidates from DH4_athlete Haplotype A against DH4_athlete Haplotype B

Table S7. BLAST Top hit results of the 132 H1 candidates from ‘Buster’ against all predicted NLRs from DH4_Athlete

Table S8. dRenSeq results of previously identified SH genes in 127 cultivars including DH4_Athlete.

Figure S1: Transformed linear genomescope2 profile of 21-mers derived from ONT reads.

Figure S2: Inter-haplotype comparison of gene pair ordering across all twelve chromosomes.

Figure S3: Genome Continuity Inspector (GCI) profiles of ONT read alignments to haplotype scaffolds. ONT read coverage is represented by the green graph, regions with potential misassemblies are marked with blue (= read depth between 0 and 0.1*mean depth) or red (= read depth 0) highlights.

Figure S4: K-mer counts associated with NLRs across candidates from the assembled haplotype A and haplotype B, as identified through the association study.

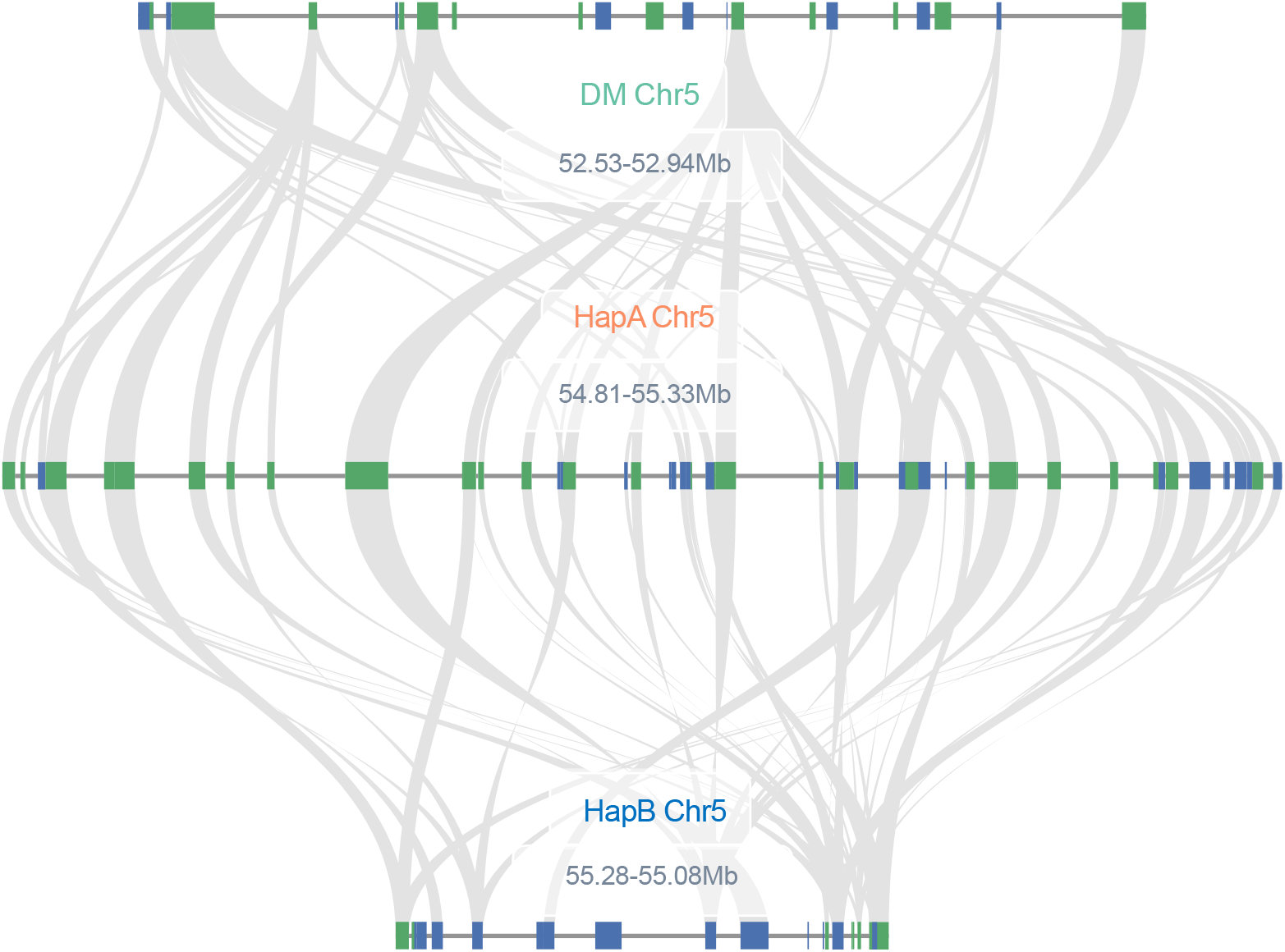

