## Supplementary figures and images for "Phased Potato Genome Assembly and Association Genetics Enable Characterisation of the Elusive H1 Resistance Locus Against Potato Cyst Nematodes"

### Figure S1

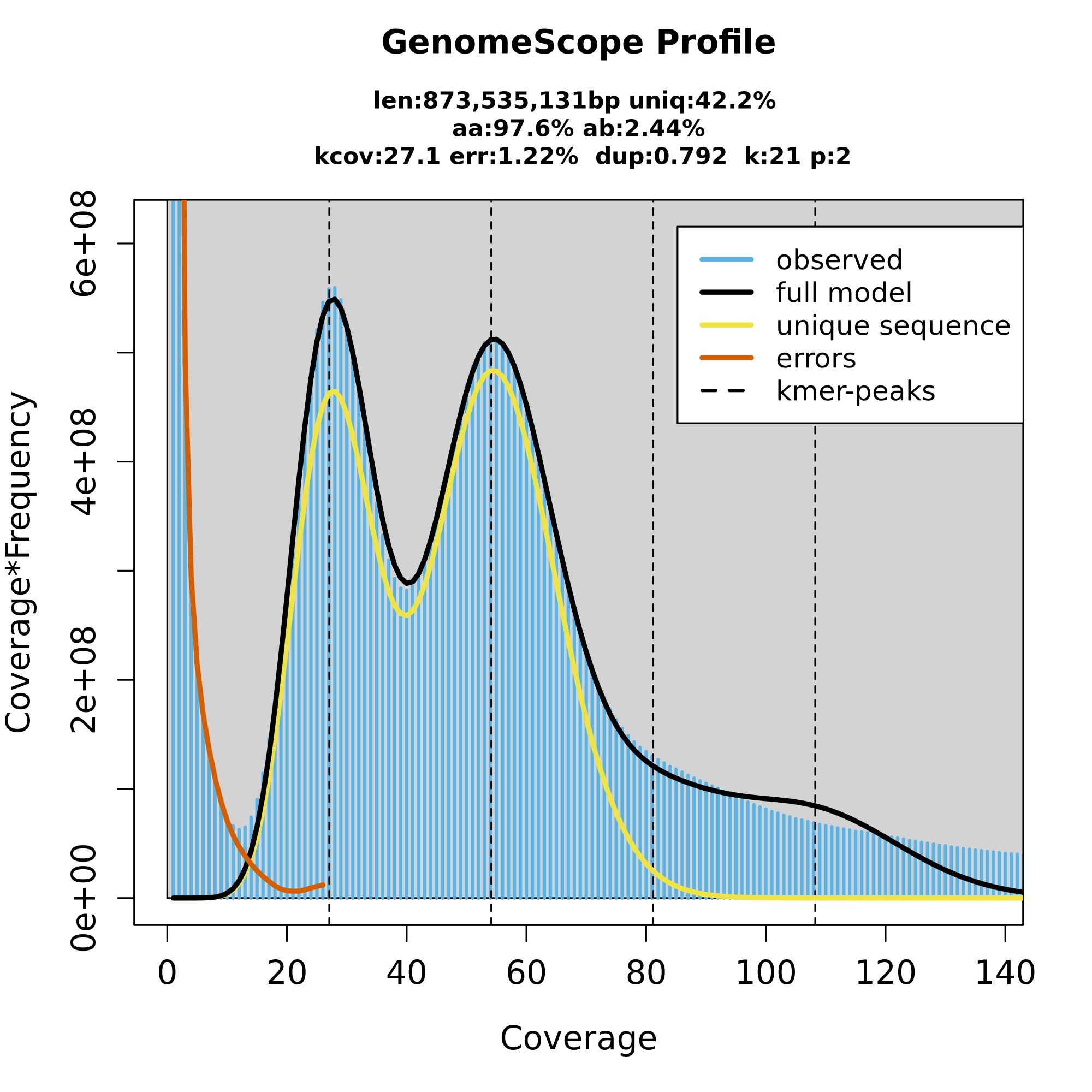

### Figure S2

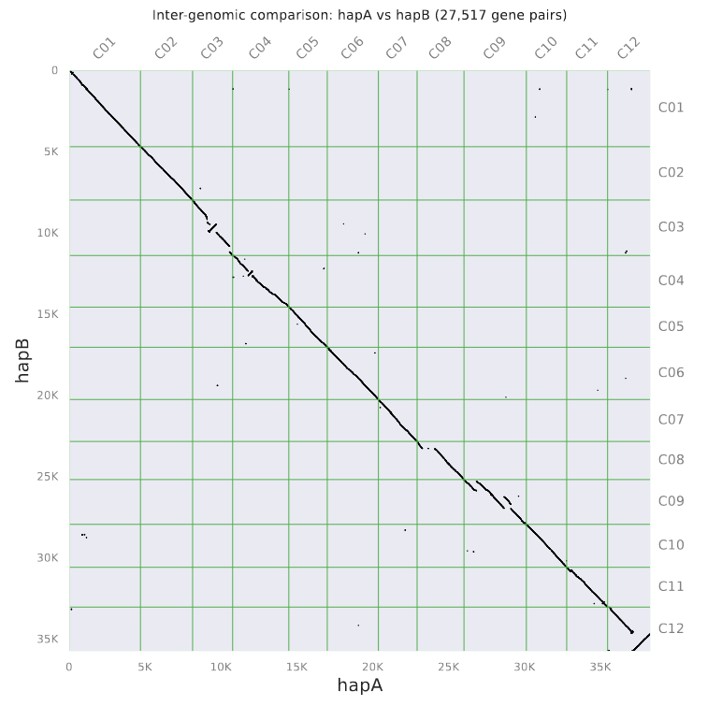

### Figure S3

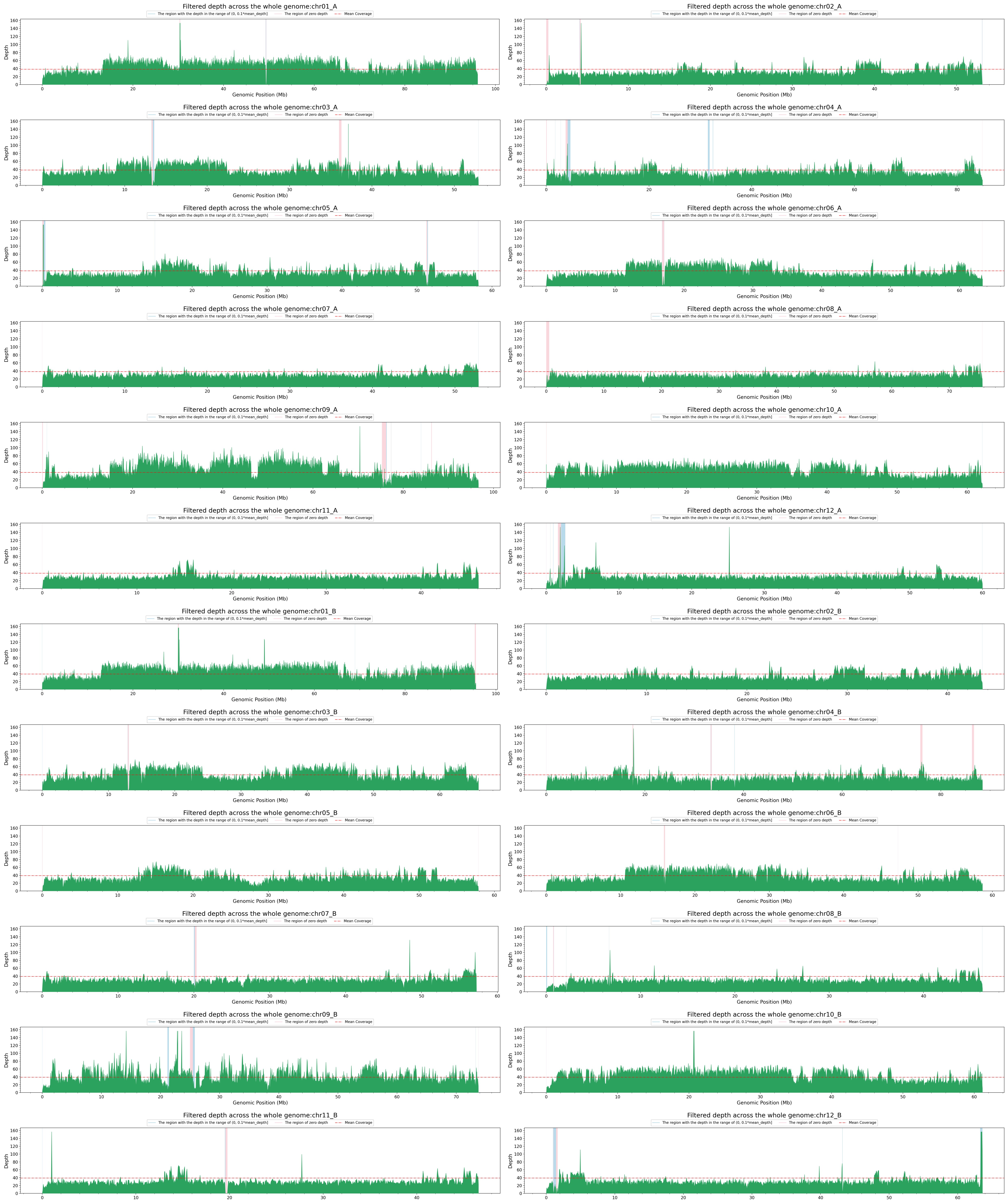

### Figure S4

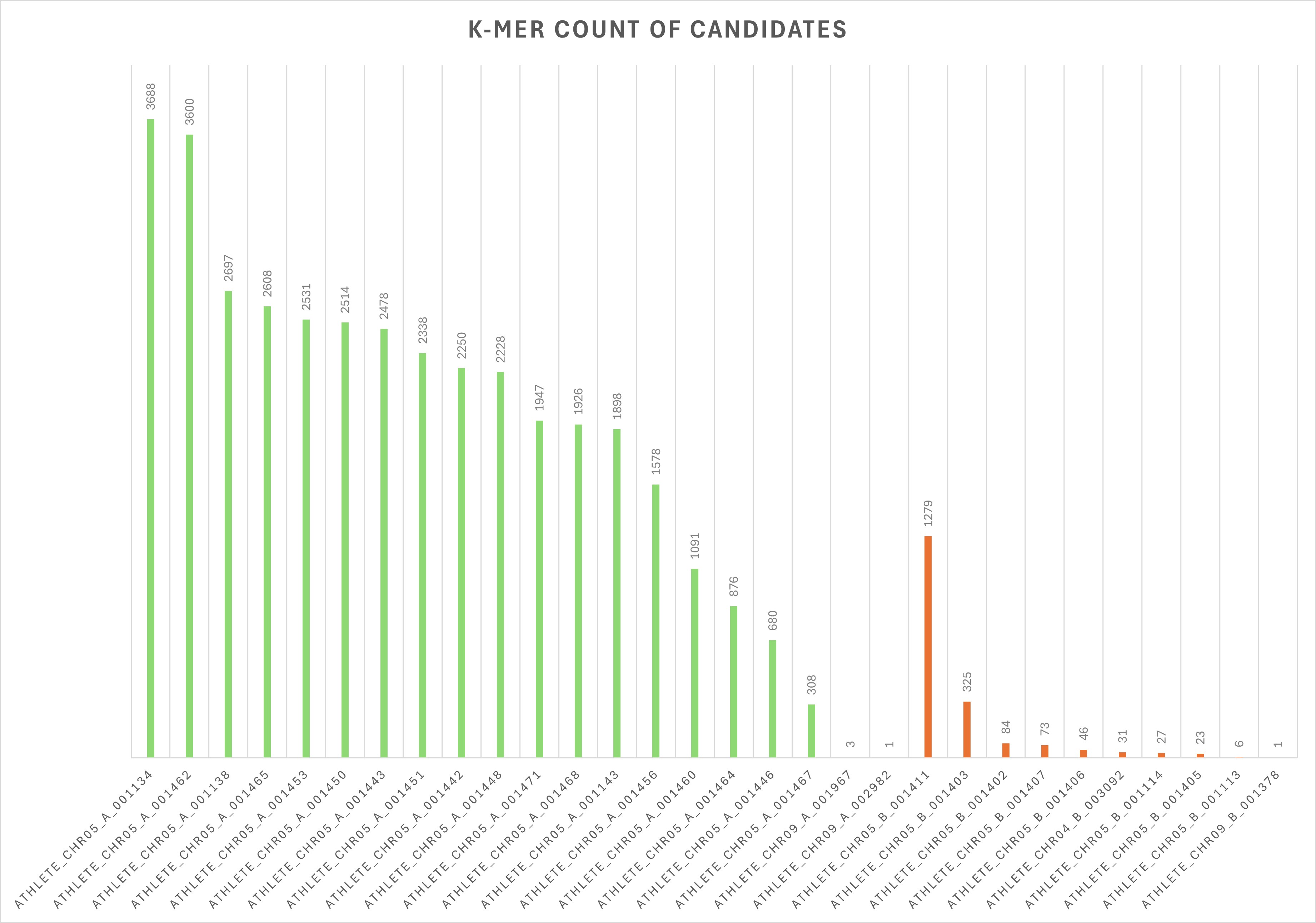
